# Deep Learning Shows Cellular Senescence Is a Barrier to Cancer Development

**DOI:** 10.1101/2021.03.18.435987

**Authors:** Indra Heckenbach, Michael Ben Ezra, Garik V Mkrtchyan, Jakob Sture Madsen, Malte Hasle Nielsen, Denise Oró, Laust Mortensen, Eric Verdin, Rudi Westendorp, Morten Scheibye-Knudsen

## Abstract

Cellular senescence is a critical component of aging and many age-related diseases, but understanding its role in human health is challenging in part due to the lack of exclusive or universal markers. Using neural networks, we achieve high accuracy in predicting senescence state and type from the nuclear morphology of DAPI-stained human fibroblasts, murine astrocytes and fibroblasts derived from premature aging diseases *in vitro*. After generalizing this approach, the predictor recognizes an increasing rate of senescent cells with age in H&E-stained murine liver tissue and human dermal biopsies. Evaluating corresponding medical records reveals that individuals with increased senescent cells have a significantly decreased rate of malignant neoplasms, lending support for the protective role of senescence in limiting cancer development. In sum, we introduce a novel predictor of cellular senescence and apply it to diagnostic medical images, indicating cancer occurs more frequently for those with a lower rate of senescence.

## Introduction

Cellular senescence is widely recognized as a fundamental process in aging, both as a primary causal factor in the decline of tissue homeostasis and as a consequence of other aging processes such as inflammation and DNA damage^1–3^. Due to its critical role in disease etiology, senescence is increasingly recognized as a target for pharmaceutical intervention^4^. It also serves as a biomarker for aging^5^, possibly providing a more nuanced measure of age-related health in model organisms beyond simple chronological age. However, the role of senescence in human health is not clearly understood. Senescent cells present a complex and diverse phenotype, which varies significantly by cell type and source^6,7^. There is considerable overlap between molecular factors that associate with senescence, DNA damage repair, inflammation, and other processes^8^. Some of the most common markers of senescence are beta-galactosidase, produced by increased expression from lysosome activity, and the cell cycle inhibitors p16 and p21. Nevertheless, there is no single marker that reliably and consistently identifies senescence^9–11^. Importantly, senescent cells often exhibit an altered morphology, including expanded nuclei and an irregular, flattened appearance^12,13^, making senescence amenable to analysis with computer vision and machine learning methods^14^.

We present deep learning models that can predict cellular senescence with high accuracy based on nuclear morphology. These methods can further distinguish between multiple types of senescence, including radiation-induced damage and replicative exhaustion. Notably, predicted senescence correlates substantially with DNA damage markers γH2AX and 53BP1 foci counts. Our senescence predictor was developed using normal human fibroblast lines, but it also identifies increased senescence when applied to multiple types of premature aging diseases, including Hutchinson-Gilford progeria syndrome (HGPS), ataxia telangiectasia (AT), and Cockayne syndrome (CS). We also evaluated the predictor on mouse astrocytes and found it indicated increased senescence in cells subjected to ionizing radiation, confirming its relevance to different cell types and organisms. These methods were applied to H&E-stained mouse liver tissue, where we found an increasing rate of senescence with age. Further, these methods were applied to H&E-stained human tissue sections and predict an age-dependent increase in senescence. Using the National Patient Register, which records all ambulatory and in-patient contacts with Danish hospitals, we investigated how predicted senescence relates to human disease. We found a highly statistically significant relationship between malignant neoplasm incidence and fewer predicted senescent cells, which fits the hypothesis that senescence is a mechanism to limit cancer^15,16^. In our study of 169 individuals, we found that a predicted senescent cell load above the age-dependent average correlated with reduced incidence of malignant skin-diagnosis at 33.3%, compared to 48.9% for patients with predicted senescence below average. Further, a predicted senescent cell load above the age-dependent average correlated with reduced incidence of non-skin related cancer at 16.0% compared to 29.5% for patients with predicted senescence below average. While oncogenic events are associated with the formation of senescent cells^15^, we speculate that individuals with higher propensity toward developing senescent cells have reduced formation of malignant neoplasm and are at lower risk of cancer.

## Results

Several fibroblast cell lines, maintained in cell culture, were treated to induce senescence by ionizing radiation (IR) or passaged until they reach replicative senescence (RS) (**Fig. S1a, b, c**). After fluorescent staining with DAPI to highlight the nuclear DNA, the cells were imaged with a high content microscope. Nuclei were detected using a deep convolutional neural network based on U-net, which produced output images containing the detected nuclear regions. Each detected nucleus was extracted into a cutout for subsequent analysis. We applied several methods to normalize features in images, such as removing the background, standardizing the size of the nuclei, and even masking inner details of the nuclei (**Fig. 1a, b**).

**Figure 1.**
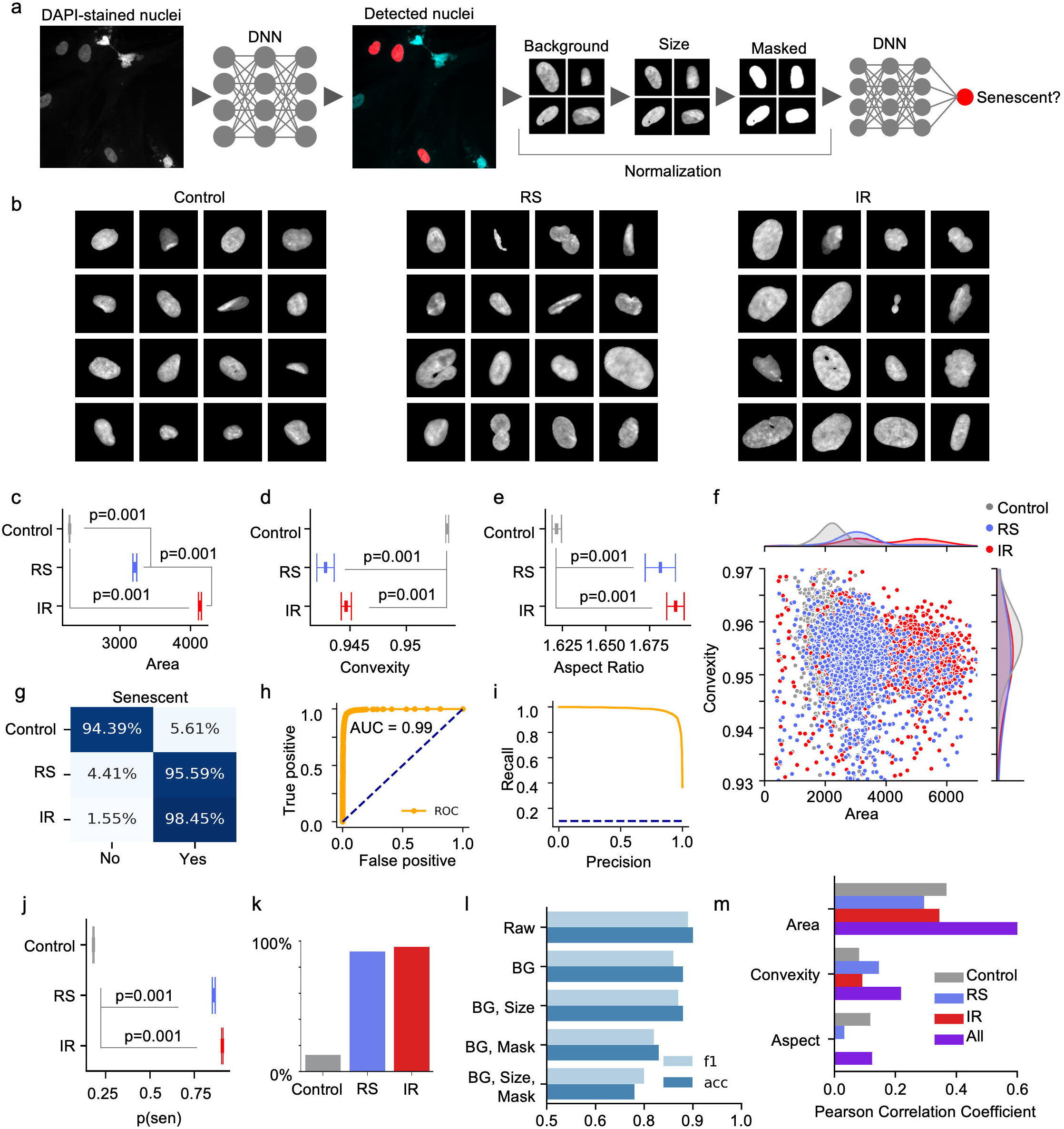
Nuclear morphology is an accurate senescence predictor *in vitro*. **a** Analysis workflow. **b** Sample nuclei for controls, replicative senescence (RS) and ionizing radiation (IR) induced senescent cells. **c** Area of identified nuclei (n=6,976-68,971, mean ± 95% CI). **d** Convexity of identified nuclei (n= 6,976-68,971, mean ± 95% CI). **e** Aspect ratio of identified nuclei (n= 6,976-68,971, mean ± 95% CI). **f** Scatter plot of individual nuclei, with overall distributions for each to the top and right. **g** Accuracy of a deep neural network (DNN) predictor on validation data. **h** Receiver operating characteristics (ROC) curve of the DNN. **i** Precision/recall curve. **j** Predicted senescence probability of nuclei for independent cell lines (n= 2,504-22,481, mean ± 95% CI). **k** Percent of nuclei in each state classified as senescent for independent cell lines. **l** Accuracy of DNNs trained and predicting after different normalization methods. **m** Correlation between morphological metrics and predicted senescence by class, BG: background.

### Senescent Cells Display Altered Nuclear Morphology

A morphological analysis of the detected nuclei was performed to compare control cells to those that were senescent. There was a significant difference in nuclear area for each of the three groups with increased nuclear area as previously reported^12^. In addition, IR senescent cells were significantly larger than RS cells (**Fig. 1c**). Aging and certain premature aging diseases have been associated with greater irregularities or folds in the nuclear envelope^17,18^. We therefore evaluated convexity, which is a ratio of nuclear area to convex hull area, as a measure of the nuclear envelope regularity. Convexity showed the shape of control cells were more regular compared to both IR and RS, which had a more irregular boundary (**Fig. 1d**). RS has the lowest convexity value, indicating the highest irregularity (or lowest regularity). This indicates convexity is another measure of senescence, with lower values corresponding to increased senescence. In addition, we looked at aspect ratio, a measure of width compared to height (measured as the longest compared to shortest dimensions of a minimized rectangle around each nucleus) and found that both IR and RS had higher values compared to controls (**Fig. 1e**). We compared area and convexity per nuclei, observing overlapping clusters for the three states with area of RS overlapping both control and IR, and convexity of RS and IR overlapping (**Fig. 1f**). Interestingly, the distribution of the area of the IR senescent cells was bimodal, with the lower mode matching RS and a higher mode at almost twice the area of the RS, perhaps suggesting IR induced aneuploidy or stalling at the G2 checkpoint of the cell cycle (**Fig. 1f**, upper histogram distribution of joint scatter plot). Simple nuclear morphological measures appear to be a viable method for assessing cellular senescence *in vitro*.

### A Deep Learning Classifier Accurately Predicts Senescence Based on DAPI staining

Given the rich structure of nuclei and potentially broad set of features, we applied deep neural networks to better assess senescence. A custom convolutional neural network was trained using 80% of the samples, while 20% was held out for validation. After seeing accuracy converge to a steady level, the model was applied to validation data. We also compared our custom network to Xception, one of the top performing models for image classification that has been often applied to biomedical classification^19,20^. Xception achieved superior results with an f1-score of 94%, accuracy of 95%, and AUC of 0.99 with validation data (**Fig. 1g, h, i**). To eliminate any potential overfitting on the experimental context and cell lines, we evaluated the model on an independent data set of two additional cell lines, which were prepared and imaged separately. This achieved an f1-score of 92%, accuracy of 94%, and AUC of 0.96 (**Fig. S2a, b, c**). The mean probability of senescence per nuclei is 0.18 for controls, 0.86 for RS, and 0.91 for IR (**Fig. 1j**), indicating senescence for 12.7% of controls, 92.0% of RS, and 95.6% for IR using the standard threshold (**Fig. 1k)**.

In another experiment, deep neural networks were trained to detect control compared to different senescent types, IR and RS. Xception, trained similar to the dual state experiment above, produced a mean class accuracy of 78.6% in detection of the three states, with 83.3% for controls, 75.7% for RS, and 76.8% for IR (**Fig. S2d, e**). It achieved a relatively high AUC of 0.9 for RS and 0.95 for IR. In sum, nuclear morphology represents a strong predictor of both replicative and DNA damage induced senescence.

### Nuclear Shape Is A Central Predictive Feature in Senescent Cells

Nuclei images contain several features that could be used for classification; however, it is unclear what the deep neural network is using as its basis for assessment. Nuclear area, staining intensity and even the image background itself could contain a signal that the neural network is picking up on. To provide some insight into how much these potential factors contribute to senescence classification, we trained several models based on reduced forms of the cutout library. Our base model already includes brightness standardization. First, the background of the nuclei was masked, by excluding all areas outside of the U-Net detected nuclear region. Next, we applied size normalization, such that the greater of the width and height was set to a standard pixel size. Finally, we converted the interior of nuclei to a single-color value, essentially masking all internal structure. With each reduction, we observed a slight decrease in classification accuracy when applied to independent test lines (**Fig. 1l**). The background masking produced 86% for the f1-score and 88% for accuracy, a small reduction indicating limited reliance of the background. With background masked and size normalized, a trained model produced 87% for f1-score and 88% for accuracy, showing area and size played little role in senescent detection. This model was further reduced by completely masking the internal structure of the nuclei, which led to an f1-score of 80% and accuracy of 78% (**Fig. S2f, g**). While masking was a significant reduction in accuracy, it is remarkable that so much information could be removed from nuclear images and still obtain a relatively accurate classification of senescence. These experiments suggest that classification is largely based on the overall shape of the nuclei. We explored this further by evaluating Pearson correlation between predicted senescence and several morphological metrics, finding that area was moderately correlated but both convexity and aspect ratio were weaker (**Fig. 1m**). The deep learning model appears to be picking up on the nuclear shape in a more sophisticated manner than simply aspect ratio or convexity.

The final reduced model yields an overall accuracy of 78%, and it shows an imbalanced per class accuracy of 73.9% for control, 69.3% for RS, and 91.4% for IR. It maintains a good AUC of 0.88. With similar reductions, the three-state senescent type detector model shows overall accuracy of only 58% (**Fig. S2h, i**). The RS state has poor accuracy at 31.3%, but 87.7% for controls and 56.1% for IR. The AUC has declined to 0.71 for RS and 0.6 for IR. Despite lowering accuracy, the feature standardization and reduction makes the model less influenced by a large number of technical variations such as image intensity, choice of staining method, magnification and others that could impact the utility of the predictor.

### Classification with Confidence

While overall accuracy per-nuclei was relatively high, a sizable number of nuclear images were ambiguous, which can be interpreted as the model being uncertain in its prediction. Extending neural networks with Bayesian properties has several advantages, most notably providing a measure of confidence for predictions^21^. The Bayesian Neural Networks (BNN) allows for the construction of a posterior probability distribution which can be used for interval estimation, compared to a single prediction from a classic neural network. Samples can be filtered to reduce the ambiguous cases by requiring higher mean probability from the BNN. Using Tensorflow Probability, we developed several BNNs. Our custom model converted to a BNN performed adequately for raw cutouts, but it would not train well for the masked/normalized nuclei. We partially converted Xception to utilize Flipout nodes^22^, leaving the separable convolutions as point estimate nodes. We also fully converted InceptionV3 as an alternative model. Our partial BNN of Xception produced an f1-score of 84%, accuracy of 86%, and AUC of 0.92 (**Fig. S2j, k**). The full BNN for InceptionV3 gave an f1-score of 79%, accuracy of 80%, and AUC of 0.87 (**Fig. S2l, m)**. The BNN models can be thus be used to understand the probability distribution of the data but at a lower accuracy.

### A Deep Neural Network Ensemble Increases Predictive Power

After training the senescent classifier through different sessions, we saw variance in the predictions for a subset of samples. Exploring a large multidimensional solution space during training, neural networks select a relatively good solution that is often biased to favor certain classes^23^. Using an ensemble of deep models, the predictions can be combined as though consulting a collection of experts (or interpreted as the “wisdom of the crowd”). To achieve this, we trained an ensemble with random initial weights, potentially allowing convergence to different local minima. We found that there is consistent agreement for the majority of samples, however, there is a significant percent of edge cases with a high variance in predictions among the model instances (**Fig. 2a**).

**Figure 2.**
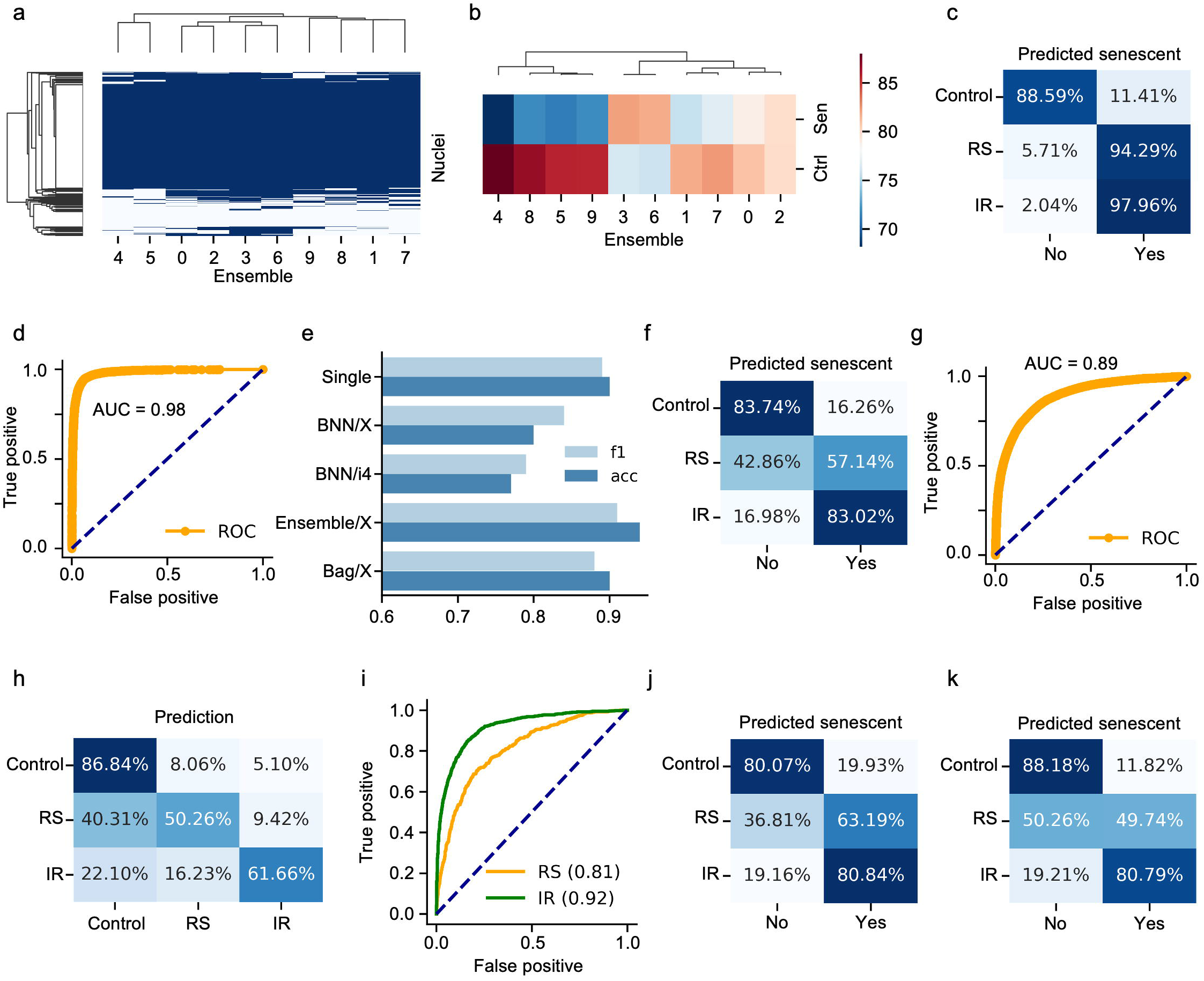
Predictions from deep ensembles. **a** Heatmap of variation in predictions by members of ensemble (500 sample nuclei as rows, ensemble members as columns). Blue is young/control and white is senescent. **b** Heatmap of per-class accuracy for control and senescent by ensemble model. **c** Accuracy of deep ensemble. **d** ROC curve for the deep ensemble. **e** Accuracy of single model, Bayesian neural networks, deep ensemble, and bagging. **f** Accuracy of deep ensemble with normalized samples. **g** ROC curve for the deep ensemble with normalized samples. **h** Accuracy of three-state senescence ensemble model. **i** ROC curve for the type ensemble model. **j** Accuracy of RS-only model. **k** Accuracy of IR-only model.

We therefore speculate that using an ensemble of deep models for inference and aggregating the results provides predictions with less bias and higher confidence (**Fig. 2b**). Evidently, some models balance the accuracy of each class in the middle of the range (75-80%), while other models skew toward one class at the expense of the other (for example, obtaining ∼85% on one but ∼70% on the other). While ensembles have benefits like a BNN, they can be less biased since each ensemble member can specialize around a solution, while a BNN is confined to a single local minima in solution space. Accordingly, we obtained good results with the ensemble method, with an f1-score of 91%, accuracy of 94%, AUC of 0.98 (**Fig. 2c, d**). More importantly, the ensemble provides a higher confidence and less biased approach by combining multiple models that specialize in predicting different classes.

### An ensemble of neural networks outperforms Bayesian neural networks

We also tried Bagging, where bootstrapping with replacement selects a subset of the samples to use in training independent models. This method did not provide a significant improvement over the basic deep ensemble method (**Fig. 2e**). The BNN models can be used to improve confidence but sacrifice performance, while the ensemble models provide both (**Fig. 2e**). We therefore further evaluated the deep ensemble method with masked and normalized samples. This produced an f1-score of 80%, accuracy of 82%, and AUC of 0.89 (**Fig. 2f, g**), which improved upon the single model. The ensemble method was also applied to the tri-state model to distinguish senescent type, which achieved overall accuracy of 66% and AUC of 0.81 for RS and 0.92 for IR (**Fig. 2h, i**). While this is lower accuracy, it is an overall improvement of 23.64% compared to the single normalized tri-state model. With all states well above the 33.3% accuracy expected from random predictions, this model is capable of recognizing type of senescence given an adequate sample size.

Due to the lower performance of senescent type prediction, we trained deep models on each type of senescence exclusively, training for control vs RS-only or control vs IR-only. This left the other state undefined, assessing each type of senescence separately. Both models classified IR with high accuracy, but the RS-only model recognized RS with ∼13% higher accuracy, while the IR-only misclassified those as control (**Fig. 2j, k**). Ensembles of deep neural networks clearly allow for greater accuracy for senescence prediction.

### Modifying Thresholds Increases the Accuracy of Prediction and Improves Confidence

Deep neural networks utilizing one-hot node outputs with the softmax function are trained to produce numerical values that are sometimes treated as the probability for each state. They should not be interpreted as model confidence, but by sampling from a BNN or deep ensemble, we can utilize the distribution to determine uncertainty^21^. We evaluated the predictions for the BNN and deep ensemble (**Fig. S3a, b**). Correct predictions are indeed oriented toward the lower and higher range of the softmax output, representing greater certainty about a sample’s state. In both cases, the incorrect predictions are clustered toward the center with the 0.5 threshold. Different models could be biased toward either state by shifting those ambiguous samples across the threshold.

We can assume higher confidence in a model’s predictions by raising the classification threshold (of both one-hot states, thereby filtering the predictions in the middle). We therefore evaluated the accuracy using a range of thresholds from 0.5 up to 0.95 in the single model, the Xception BNN, the ensemble of models, and the ensemble of fully normalized models (**Fig. S3c, d, e, f)**. In all cases, we see a significant increase in accuracy as the threshold is raised, due to the ambiguous samples being discarded. By raising the threshold, the Xception-based BNN goes from 85.6% to 96.0%, while the ensemble of normalized models goes from 81.6% accuracy to 97.2%. A similar approach was applied to other models, including the IR-only and RS-only models (**Fig. S3g, h**). Raising the threshold, these also showed a gain in accuracy of 10-15%. Unfortunately, this led to a significant reduction in the number of samples considered. There is a tradeoff between number of predictions and accuracy, which must be balanced for each application to ensure suitable power for analysis.

The tri-state model, which distinguishes between IR and RS, showed lower accuracy, especially when applied to the fully normalized samples (**Fig. S2h, i**). As a deep ensemble, we see accuracy of 86.8% for control, 50.3% for RS, and 61.7% for IR. Since there are three states, even the 50.3% accuracy with RS places the majority of its samples in the correct category, with 40.3% a FN appearing as control and 9.4% as IR. It’s ROC curve has AUC of 0.81 and 0.92 for IR. Applying threshold adjustments, we see the overall accuracy go from 80% up to 95%. Maintaining a majority of samples, a threshold of 0.8 exceeds 90% accuracy.

### DNA Damage Foci and Area Correlates with Senescent Prediction

Senescent cells are associated with permanent increase in nuclear foci of the DNA damage markers γH2AX and 53BP1^24,25^. We characterized the DNA damage response (DDR) foci for our cell lines and investigated how these foci relate to predicted senescence. Our base data set including control, RS, and IR lines were examined for damage foci. Using high content microscopy, we counted DNA damage foci per nuclei and found the mean count of γH2AX and 53BP1 foci to be below 1 each (0.9 and 0.6, respectively) for controls, while RS had 4.0 γH2AX and 2.0 53BP1 foci and IR had 3.4 Hγ2AX and 3.0 53BP1 foci (**Fig. 3a, b, S4a**). To study how the presence of damage foci relates to predicted senescence we calculated the Pearson Correlation between predicted senescence and γH2AX and 53BP1 foci counts. We found that across all conditions there is a moderately strong correlation of around 0.5 (**Fig. 3c)**. This association is also visible when simply plotting foci counts and senescence prediction which shows predicted senescence flipping from low to high, along with shifts in foci counts (**Fig. S4b**). The same pattern applies to area, with shifts in the concentration of area along with shifts in the predicted senescence, aligning well with cell conditions. Within senescent subtypes RS and IR, the correlation is slightly weaker, perhaps indicating that the senescent probability score for each subtype has some correlation with foci count. Our feature reduction including masking means that internal nuclear structure was not used in assessment, but it is nonetheless notable that senescence prediction (overall and by subtype) correlates with foci count. We also compared the correlation between predicted senescence and area. Here too, we see a correlation of around 0.5, and slightly weaker for the subtypes. In sum, there is a considerable correlation between foci counts and senescence.

**Figure 3.**
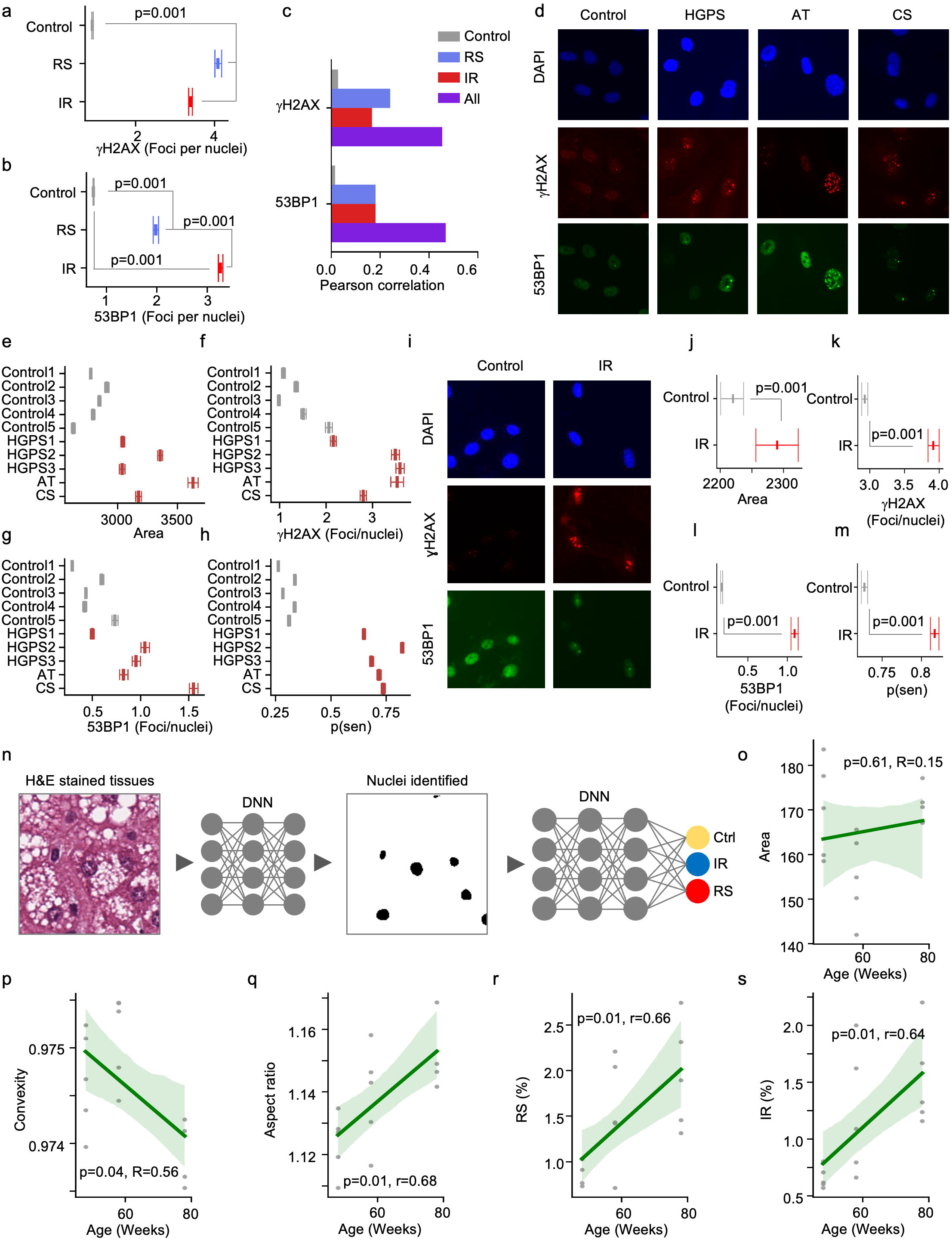
Senescence can be predicted across tissues and species. **a** Number of γ H2AX foci by type of senescence (n=1,831-15,560, mean ± 95% CI). **b** Number of 53BP1 foci by type of senescence (n= 1,831-15,560, mean ± 95% CI). **c** Correlation between foci count and predicted senescence. **d** Representative immunohistochemistry micrographs of premature aging nuclei with DNA damage foci staining. **e** Nuclear area for premature aging diseases (n=4,340-15074, mean ± 95% CI), HGPS: Hutchinson-Gilford Progeria Syndrome, AT: ataxia telangiectasia, CS: Cockayne Syndrom. **f** Number of gH2AX foci for premature aging diseases (n= 5,162-17,584, mean ± 95% CI). **g** Number of 53BP1 foci by premature aging diseases (n= 5,162-17,584, mean ± 95% CI). **h** Predicted probability of senescence for premature aging disease (n=5,162-17,584, mean ± 95% CI). **i** Representative immunohistochemistry micrographs of senescent murine astrocytes with DNA damage foci staining. **j** Area of murine astrocytes (n=4,888-13,549, mean ± 95% CI). **k** Number of gH2AX foci for murine astrocytes (n=4,918-13,661, mean ± 95% CI). **l** Number of 53BP1 foci for murine astrocytes (n= 4,918-13,661, mean ± 95% CI). **m** Predicted senescence for murine astrocytes (n= 4,918-13,661, mean ± 95% CI). **n** Analysis workflow. **o** Mean nuclear area per mouse by age (n=5). **p** Mean nuclear convexity per mouse by age (n=5). **q** Mean nuclear aspect ratio per mouse by age (n=5). **r** Predicted percent that are RS senescent (n=5). **s** Predicted percent that are IR senescent (n=5).

### Progeria Cell Lines Display Increased Senescence

Patients suffering from premature aging, or progeria, represent genetically well-defined models to understand the molecular basis of aging^26,27^. To test if cell lines from progeria patients display accelerated aging *in vitro*, we applied the senescent classifier to primary fibroblasts isolated from Hutchinson-Gilford progeria syndrome (HGPS), ataxia telangiectasia (AT) and Cockayne syndrome (CS) (**Fig. 3d**). Evaluating the area of the nuclei of progeria cells, we found that in general their mean is significantly larger than controls. Notably ataxia-telangiectasia cells have the largest nuclei at 25% higher than controls, while Hutchinson-Gilford progeria and Cockayne syndrome are both 15% higher (**Fig. 3e**). We also investigated DNA damage foci and observe that most prematurely aged lines have higher γH2AX and 53BP1 foci counts (**Fig. 3f, g** and **Fig. S4c**). Further, despite diverse mechanisms, the classifier recognized these cell lines having significantly greater probability of senescence **(Fig. 3h)**. All progeria lines have high mean probability of senescence at 0.7, indicating that the average cell in each group is considered senescent, while controls are below the standard threshold at 0.3. These observations suggest that our classifier may be able to discriminate rates of aging *in vitro*.

### The senescent classifier translates across species and cell types

To broaden the applicability of our classifier we speculated that it might apply to nuclei from other cell lines and species. We therefore evaluated the model on mouse astrocytes, which were treated with IR (**Fig. 3i**). We first compared the nuclei area and found that the IR treated astrocytes had slightly but significantly larger nuclei than controls (**Fig. 3j**). To test if senescence classification is based on area, we calculated the Pearson Correlation Coefficient between these two measures. With a PCC=0.12 and p-value 4.6×10^−69^, we find only a weak relationship between area and senescence. Evaluating DNA damage foci, we see that IR treated astrocytes have substantially higher foci count as expected (**Fig. 3k, l, and S4d**). We next applied the ensemble of deep models (without normalization) and found that the IR treated cells had a 9% higher probability of senescence than controls (**Fig. 3m**).

We also applied the model to H&E stained liver tissue from C57Bl6 mice at taken at 48, 58, and 78 weeks of age. After imaging the tissue sections at 20x, we used a deep learning segmentation model trained on 18 tiles to extract nuclei from 16,187 tiles (**Fig. 3n**). We first analyzed morphological metrics, finding an insignificant increase in nuclear area (**Fig. 3o**). However, we saw a significant decrease in convexity and increase in aspect ratio, both indicating increased senescence with age (**Fig. 3p, q**). Nuclei were evaluated for senescence using the normalized RS-only and IR-only models, of which the RS model indicated increasing senescence with age while the IR model did not significantly increase (**Fig. S4e, f**). Using the probability, we calculated the percent of senescent cells, finding ∼36% for RS and ∼99% for IR. The predictor is trained on *in vitro* DAPI stained fibroblasts representing a considerable difference in context, it is therefore likely that the algorithm should be tuned to evaluate other data sources. Applying thresholds of 0.8 and 0.94 for RS and IR, respectively, the percent was brought down to roughly 1-2% each to match the reported senescent rate in the liver^28^. With these thresholds, the percent of senescent cells per mouse increased with age (**Fig. 3r**,**s**). Given the differences in human and mouse nuclei as well as between cell types, it is notable that the senescent state can be captured through the relative difference in assessed probability. It therefore appears that our predictor may be able to determine senescence across cell types and species.

### The human dermis shows age-dependent increase in senescent nuclei

To further investigate if the predictor could be applied in a clinical context, we tested the algorithm on human skin samples of 169 individuals aged 20-86 years. The senescent classifier was evaluated on the dermal nuclei from biopsy samples stained with H&E and imaged in a slide scanner at 20x. We applied U-Net to detect nuclei, extracted nuclear regions, and converted the nuclei to the normalized and masked form (**Fig. 4a**). We first evaluated several morphological metrics, including area, convexity, and aspect ratio. Across age, we see no change in area (**Fig. 4b**), an insignificant change in convexity (**Fig. 4c**), and a significant change in aspect ratio (**Fig. 4d**). Applying the senescent predictor, the probability of senescence increases with age of patients for RS but is relatively flat for IR (**Fig. S5a, b)**. We applied the standard softmax threshold and evaluated the percent of cells considered senescent, which yielded means of 25% for RS and 40% for IR. The percent was significantly higher than we’d expect for human dermal nuclei, ranging from mean of ∼1% in young to ∼15% in old^28^. Our fully normalized model has a relatively high FP rate (20% for RS and 12% for IR), and human dermal nuclei are disproportionately non-senescent, likely exaggerating the predicted senescence. We therefore adjusted the threshold to reduce false positives and attempt to compensate for the large biological difference. To calibrate the model to the level of senescence expected for dermal nuclei, we set the cutoff to 0.7 for RS and 0.85 for IR, which lowered the percent for all patients to a mean of ∼6% each and showed an age-dependent increase in percent of senescence (**Fig. 4e, f**). We also evaluated the correlation between morphological metrics and predicted senescence and found moderate correlation for several metrics, but RS was more correlated with convexity while IR was more correlated with area and aspect ratio, perhaps indicating morphological aspects of each type of senescence *in vivo* (**Fig. S5c**). Interestingly, we found that area was anti-correlated with both forms of predicted senescence and predicted probability of IR was inverse to aspect ratio (**Fig. S5c, d)**. This again emphasizes the difference between senescence *in vitro* and in tissue and also affirms that the IR and RS model are picking up on different aspects of senescence.

**Figure 4.**
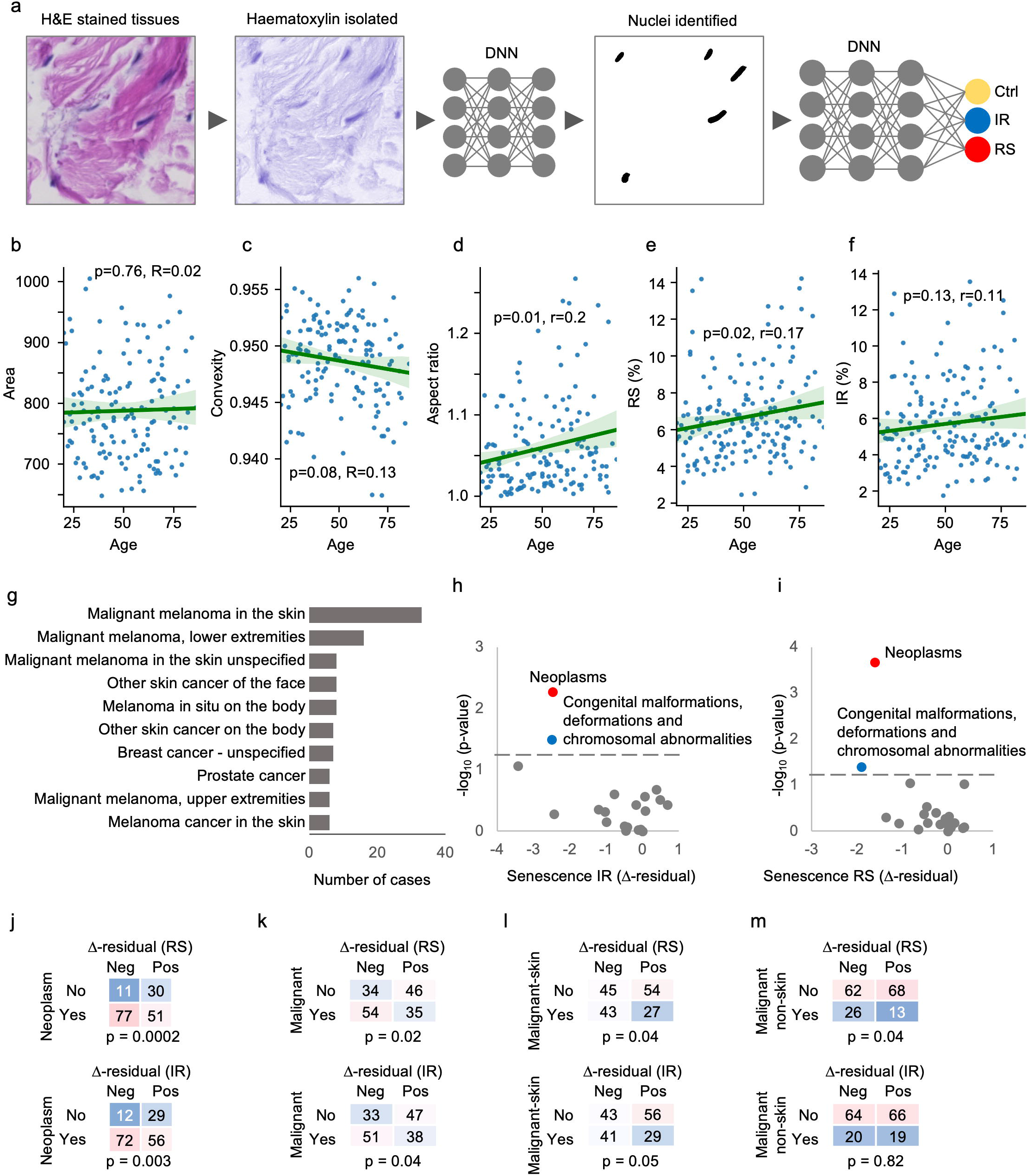
Nuclear morphology predict senescence and cancer risk in humans. **a** Analysis workflow. **b** Mean nuclear area per patient by age (n=148). **c** Mean nuclear convexity per patient by age (n=148). **d** Mean nuclear aspect ratio per patient by age (n=148). **e** Predicted percent that are RS senescent (n=169). **f** Predicted percent that are IR senescent (n=169). **g** Number of cases for most common cancer conditions. **h** Volcano plot of conditions based on IR senescence residuals and chi-square p-values. **i** Volcano plot of conditions based on RS senescence residuals and chi-square p-values. **j** Contingency table between neoplasms and residuals of predicted senescence. **k** Contingency table between malignant skin neoplasms and residuals of predicted senescence. **l** Contingency table between all malignant neoplasms and residuals of predicted senescence. **m** Contingency table between malignant non-skin neoplasms and residuals of predicted senescence.

### Senescent dermal nuclei are inversely associated with neoplasms

Given the large variation in predicted senescence, we speculated that these values could represent meaningful health outcomes. To investigate, we retrieved ICD-10 diagnosis codes collected in the Danish National Patient Register from 1977 to 2018 for all the individuals in the study (**Fig. 4g)**. We looked for associations between individuals with diagnosed conditions and predicted senescence above or below the age-dependent mean (those above or below the trendline in **Fig. S5a, b**, specifically using residuals from linear regression of RS versus age or IR versus age), using the chi-square test for the frequency of occurrence between the two groups (**Fig. 4h, i**). Remarkably, we found a significant correlation between a rate of senescence below the age-matched mean and the presence of ICD-10 Chapter II Neoplasm diagnosis codes for both RS and IR, with p-values of 0.0002 and 0.002, respectively (**Fig. 4j**). Narrowing down the analysis we determined the association was based on malignant (versus benign or unknown) codes within ICD-10 Chapter II Neoplasm with IR p-value at 0.037 and RS at 0.018 (**Fig. 4k**). Furthermore, grouping specific cancer codes together, we determined that RS is significant for both skin and non-skin cancer, with p-values at 0.041 and 0.037 respectively (**Fig. 4l, m**). IR was non-significant for non-skin and on the edge of significance for skin with p-value at 0.053. Notably, RS better represents replicative senescence which occurs naturally with age, while IR better represents DNA damage, although there is considerable overlap in predictions between the two with this model. Overall, we found that high assessed senescence corresponds to fewer neoplasms and malignancies, including both skin and non-skin.

## Discussion

In this paper we present a neural network that can predict cellular senescence based on nuclear morphology. Trained on fibroblasts maintained in cell culture, the classifier achieves very accurate results, which was confirmed by applying it to independent cell lines. We also trained models to correctly distinguish between senescence caused by radiation induced damage and replicative exhaustion. By training additional models on samples with reduced features, we infer that the shape of the nucleus alone provides a significant signal to indicate senescent state.

DAPI-stained nuclei with background removed, size normalized, and internal structure masked are still classified with high accuracy. These feature reduction methods serve a secondary purpose, making a model robust to technical variation - our neural network trained on reduced samples can make predictions on nuclei that were prepared in other experimental and imaging contexts. Indeed, the predictor distinguished senescent astrocytes, predicted an age-related increase in senescent liver cells, and confirmed senescence in cell lines from patients suffering from premature aging. Although it is still debated if universal markers of senescence exist, our findings suggest that at least morphological alterations in nuclei may be common across some tissues and species.

Our data shows that individuals with a predicted higher rate of senescent cells have reduced neoplasms and malignant cancer, in comparison to those with a lower rate of senescence. This is highly consistent with the notion that senescence is a likely mechanism to control cancer development by limiting uncontrolled proliferation^29^. Further, premalignant tumors express markers of senescence, which are absent in malignancies, and malignant tumors can regress and undergo senescence by switching off oncogenes^15^, supporting the protective role of senescence in blocking the progression of neoplasms to malignancies. In addition, loss of central senescence inducers such as p16 are very common in many cancer types^30^. Of note, there is also evidence suggesting that cellular senescence promotes malignancy through the inflammatory senescence associated secretory phenotype (SASP)^31^, that senescent cells may appear in areas where tumors tend to subsequently develop^32^, and that senescent cells and SASP induced by cancer treatment led to worse survival and healthspan^33^. While the role of senescence in cancer is highly complex, our results based on clinical data support the overall protective role for senescence in human health.

We also investigated how our deep learning predictor results correspond to other measures of senescence. Nuclear area is known to expand during senescence^12,34,35^, and we confirmed this in our *in vitro* data set, with significant differences in IR and RS senescent cells. On a per nuclei basis, we found a moderate correlation between area and predicted senescence. However, due to our size normalization, it is unlikely this classic feature is the primary signal for our deep learning model (at least for the size-normalized version). We also identified convexity and aspect ratio as key morphological properties that differ between control and senescent cells *in vitro* and found moderate correlation between each of these properties and predicted senescence. Interestingly, we found no increase in area with age in the human dermis, but a significant increase in aspect ratio and significant decrease in convexity, indicating nuclei becoming stretched and irregular with advancing age in humans. These observations confirm that size normalization is necessary to generalize our neural network classifier. It also demonstrates the value of our feature-neutral approach, where the neural network is trained to identify senescence from rich image data, and it is only later reduced through feature removal.

In sum, our deep neural network model is capable of accurately predicting the senescent state and type from nuclear morphology using several imaging techniques and has been demonstrated with several diverse applications. We applied the predictor to human skin samples and observed an age dependent increase in senescence. Remarkably, individuals who appear to have higher rates of senescent cells show reduced incidence of malignant neoplasms. This supports the long-standing hypothesis that senescence is a mechanism to limit cancer.

## Methods

### Cell culture

All human-derived primary skin fibroblast cells were purchased from Coriell Institute (USA). Control fibroblasts included AG08498 (male, 1 year), GM22159 (male, 1 day), GM22222 (male 1 day), GM03349 (male, 10 years) and GM05757 (male, 7 years). Cells were cultured at 37C and 5% CO2 either in 1:1 mix of DMEM GlutaMAX (Gibco, 31966047) and F-12 media (Gibco, 31765068) for AG08498, GM22159 and GM22222 or in EMEM media (Biowest, L0415-500) for GM03349 and GM05757. Fibroblasts derived from Hutchinson-Gilford progeria syndrome patients included AG06917 (male, 3 years), AG06297 (male, 8 years) and AG11513 (female, 8 years). Fibroblasts sampled from ataxia telangiectasia and Cockayne syndrome patients were GM03395 (male, 13 years) and GM01428 (female, 8 years), correspondingly. Cells were cultured at 37C and 5% CO2 in MEM media (Lonza, BE12-662F). Freshly isolated primary mouse astrocytes were kindly provided by the Department of Drug Design and Pharmacology, University of Copenhagen. Cells were cultured at 37C and 5% CO2 in DMEM GlutaMAX (Gibco, 31966047). All used media were supplemented with 10% fetal bovine serum (Sigma-Aldrich, F9665) and 100 U/mL penicillin-streptomycin (Gibco, 15140163).

### Senescence induction

To achieve replicative senescence control fibroblasts at early passages were seeded in T25 cell culture flasks (200 000 cells) and cultured over 32 weeks. After each splitting cell number was recorded and population doubling level (PDL) was calculated as Log_2_(cell number during harvesting/cell number during seeding). Experiment was terminated when PDL reached zero. DNA damage-induced senescence was performed according to reference^36^. Briefly, control fibroblast cells at yearly passages were seeded in 96 well plates (Corning, 3340) in a density of 2 000 cells per well. Day after cells were exposed to 10Gy of ionizing radiation and cultured for the next nine days. Medium was replaced every two days. Three days before radiated cells reached senescence state, fibroblast cells from the same stock were seeded (2 000 cells/well) as mock-irradiated control.

### Immunocytochemistry, SA-bGAL detection and image preparation

For detection of persistent DNA damage foci, fibroblast cells were washed once with warm PBS, fixed in 4% paraformaldehyde (PFA) for 15 min followed by permeabilization step with incubation for 10 min in PBS-0.1% Triton X100. Blocking was performed in 1% BSA-PBS-0.1% Tween 20 overnight at 4C. Next day cells were incubated with primary antibodies (γH2AX, 1:1000, Millipore, 05-636 and 53BP1, 1:2000, Novus, NB100-304) for 1h at RT, washed three times with PBST and incubated with secondary antibodies (1:200 Alexa-Flour 488, Invitrogen, 10424752 and 1:200 Alexa-Flour-568, Invitrogen, 10348072) for 1h at RT. Cells were incubated with DAPI solution (AppliChem, A4099) for 10 min and stored in PBS at 4C until the analysis. SA-bGAL was detected using senescence cells histochemical staining kit (Sigma-Aldrich, CS0030) according to manufacturer’s protocol. Cell colonies were imaged using INcell analyzer 2200 high content microscopy at 20x magnification to produce 1199 images with 2048×2048 pixel resolution. Due to system constraints for object detection, each image was split into four tiles of 1024×1024 pixel resolution.

### Nuclei Detection

A base library was prepared using controls, irradiated (IR), and cells serially passaged until they reached senescence (replicative senescence, RS). A deep neural network model was applied to detect DAPI-stained nuclei. The samples were used to build a training set for nuclei recognition. Several images were selected arbitrarily from each group for a total of ∼20 samples, and using custom software all nuclei in the training samples were annotated by selecting the nuclear region. U-NET, a 23-layer fully convolutional network for image segmentation, was trained using the samples, learning to associate the DAPI images with annotation masks indicating nuclear regions. Our implementation of U-NET is largely based on the original U-NET^37^, but includes a dropout layer after each of the convolutional and deconvolutional layers to reduce overfitting. After training for 1000 epochs, the U-NET model was used to detect nuclei for all 4796 tiles (1199 images x 4 tiles/image), producing output images of predicted nuclei regions. The images with predicted nuclei were scanned for recognition regions of area between 500 and 15,000 pixels. Each detected nucleus was extracted along with its surrounding context as a centered 128×128 pixel region and used to assemble a base library of 95,152 nuclei. In addition, the recognition region itself was cutout, providing a two-color reduction of the detected nuclei, and assembled into a secondary library of nuclei masks.

### Nuclear Morphology

An analysis of the nuclei was performed to assess morphological properties. The cutout nuclei were analyzed using image processing methods, such as Gaussian blur and Otsu thresholding. While these methods generally performed well for DAPI-stained nuclei, it was unsuitable for related data sets (H&E-stained histology images). Instead, the two-color masks library was used, since it provided a universal representation of the detected nuclei (with U-NET detector models that have good coverage of the nuclei region). Nuclear morphology was assessed using several metrics, including area, perimeter, moments, convexity, and aspect ratio. Convexity is the ratio of perimeter to convex hull perimeter, which provides a size-neutral measure of boundary regularity. The convex hull is a polygon that connects the outer edges of nuclei like an envelope.

### Senescent Classification

After assembling a library of senescent cells, a deep neural network was trained to classify DAPI-stained nuclei as senescent or non-senescent. Training samples were randomized and split into 80% for training and 20% for validation. Due to experimental setup, the sample classes are unbalanced, with 75.2% control, 11.2% RS, and 13.6% IR. The samples were balanced during training by applying class weights with inverse proportion to the class abundance (for example, *senescent* samples composed of IR and RS were fewer in number and therefore valued 3x higher than controls). Image samples were normalized for brightness/intensity by adjusting each image’s mean intensity to 0 and standard deviation to 1. Augmentation was also applied during training, randomly modifying samples: adjusting size from 80% to 120%, changing normalized brightness from 70% to 130%, flipping horizontally and vertically, and rotating up to 180 degrees. For each epoch, one augmentation cycle was performed. Training was done with Xception, a 48-layer model, initialized with ImageNet weights but set to allow weight adjustment of all layers during training. The top layer was replaced by a layer of one-hot nodes to indicate the state as controls or *senescent* (or as a tri-state model with controls, IR, or RS to indicate the type of senescence). With this minor adjustment, the model provided 37,640,234 trainable parameters. Training was done using Adam with the learning rate set to 1×10^−4^ for 10 epochs, in which time accuracy rapidly converged to a steady level. In addition, a simpler custom model was tested, with three convolutional layers with ReLU activation and two dense layers with L1/L2 regularization of 0.05/0.05 and 30% dropout. This model required 713,296 parameters. For both network designs, we trained with raw images along with several modified image sets, where the background was removed, the nuclei were size normalized, and the inner details of nuclei were entirely masked (**Fig. 1a**). All three techniques were based on the detected nuclei. To remove the background, the area outside of the nuclei was set to 0. Size was normalized by rescaling all nuclei so the larger of the two dimensions was a standard size of 80 pixels. Finally, the size-normalized detection region was used for the masked nuclei set.

### Bayesian Neural Network

We used Tensorflow Probability to create a Bayesian neural network (BNN). We first converted the simple custom model, replacing nodes with the comparable FlipOut version^22^, which assumes that the kernel and bias are drawn from a normal distribution. During a forward pass, kernels and biases are sampled from posterior distribution. Targets were encoded as above, and the loss function used was cross entropy plus KL divergence divided by number of batches. We also partially converted Xception to a BNN by replacing all dense and convolutional layers to FlipOut nodes, leaving separable convolutions unconverted since a FlipOut version was not available. In addition, we fully converted InceptionV3 for evaluation. Inference was done by evaluating the model 20 times to produce a distribution of predictions, and then taking the mean probability for each sample.

### Deep Neural Network Ensemble

To improve accuracy and provide a more robust solution, we also worked with an ensemble of deep learning models. This method utilized 10 models of Xception, each trained on the same data set with different random weight initialization. To generate predictions, each model instance was applied, and the results combined by taking the mean prediction. We also tried bagging, also known as bootstrap aggregation. Similar to the deep ensemble, this method trains different model instances with bootstrap selection of samples for n=1-1/e. With each instance trained on a different subset of samples, this method produces multiple models that in theory can specialize to different sets of data.

### Statistical Methods

All comparisons with between groups of samples were made using one-way ANOVA f-tests to evaluate differences in the means, followed by pair-wise tests using Tukey’s HSD (Honest Significant Difference) to calculate p-values between groups. Linear regression methods were evaluated with R and p-value statistics. Groups of patients were compared using the chi-squared test to detect significant differences between frequencies. Correlation was evaluated using the Pearson colocalization coefficient.

### Pathology sample selection

The individuals were sampled from patients for whom samples of naevi on non-sun exposed skin had undergone pathology without malignant findings at a major pathology department in Copenhagen. The patient sample was selected to have flat distribution of age. We selected patient samples from the Danish National Register of Pathology requisitioned in 2007-2017 and coded with one or more PatoSNOMED topology code: T02530 (Skin on penis), T76330 (Foreskin), T80200 (Mons pubis), T02471(Skin on nates), T02480 (Skin on abdomen), T02430 (Skin on breasts) and one or more procedure code: P30620 (resect), P306X0 (ectomy preparation), P30611 (excision biopsy) and one or more morphology code: M87400 (junction naevus), M87500 (dermal naevus), M87600 (compound naevus).

### Senescence and Human Morbidity

We collected ICD-10 diagnosis codes from the Danish National Patient Register in the period 1977-2018 of each of the patients in this study. We further grouped diagnoses into each of 21 ICD-10 chapters. We calculated the linear regression residuals of the relationship between age at pathology examination and the predicted senescent cell load (IR, RS metrics) for each of the patients. We then constructed contingency tables counting the number of patients with and without a specific diagnosis and with a predicted senescent cell load above or below the age-dependent average. We used Pearson’s chi-squared test to determine whether patients with a predicted senescent cell load above or below the age-dependent average were associated with a higher or lower incidence of specific diagnosis codes (or diagnosis within a specific ICD-10 chapter.)

### Animals

Male C57BL/6J mice were acquired from Janvier Labs (Le Genest Saint Isle, France). Animals arrived at 5-8 weeks of age and were housed in a controlled environment (12 h light/dark cycle, 21 ± 2 °C, humidity 50 ± 10%). Stratification and randomization into individual diet groups were based on baseline body weight. Mice had *ad libitum* access to tap water and chow (2018 Teklad Rodent Diet, Envigo, Madison, WI, United States; Altromin 1324, Brogaarden, Hoersholm, Denmark). The study was approved by The Institutional Animal Care and Use Committee at MedImmune (Gaitherburg, MD, United States) and The Danish Animal Experiments Inspectorate (license: 2017-15-0201-01378) and performed in accordance with internationally accepted principles for the use of laboratory animals.

### Liver histology

Terminal liver samples were dissected from the left lateral lobe immediately after sacrificing the animal and subsequently fixed overnight in 4% paraformaldehyde. The liver tissue was then paraffin-embedded and sectioned at a thickness of 3 µm. Sections were stained with hematoxylin-eosin (HE, Dako, Glostrup, Denmark). Slides were scanned by ScanScope AT System (Aperio, Vista, CA, United States).

## Acknowledgements

This research was supported by the Novo Nordisk Foundation Challenge Programme (#NNF17OC0027812), the Nordea Foundation (#02-2017-1749), the Neye Foundation, the Lundbeck Foundation (#R324-2019-1492), the Ministry of Higher Education and Science (#0238-00003B) and Insilico Medicine.

## Contributions

I.H. wrote the article, developed and trained deep learning models, and analyzed data. M.B.E. analyzed clinical data. G.V.M. performed experiments on the base data set, astrocytes, and premature aging disease. J.S.M. developed Bayesian networks and advised the project. M.H.N. and D.O. performed animal experiments. L.M. managed clinical images and medical records. E.V. advised and edited the project. R.W. advised and edited the project. M.S.K. conceived the idea, supervised the project and edited the manuscript.

## References

1. Kirkland, J. L. & Tchkonia, T. Cellular Senescence: A Translational Perspective. EBioMedicine 21, 21–28 (2017).

2. Zglinicki, T. von, Saretzki, G., Ladhoff, J., Fagagna F. d’Adda di & Jackson, S. P. Human cell senescence as a DNA damage response. Mech. Ageing Dev. 126, 111–117 (2005).

3. Covarrubias, A. J. et al. Senescent cells promote tissue NAD+ decline during ageing via the activation of CD38+ macrophages. Nat. Metab. 2, 1265–1283 (2020).

4. Childs, B. G. et al. Senescent cells: an emerging target for diseases of ageing. Nat. Rev. Drug Discov. 16, 718–735 (2017).

5. Schafer, M. J. et al. The senescence-associated secretome as an indicator of age and medical risk. JCI Insight 5, e133668 (2020).

6. Young, A. R. J., Narita, M. & Narita, M. Cell Senescence as Both a Dynamic and a Static Phenotype. in Cell Senescence (eds. Galluzzi, L., Vitale, I., Kepp, O. & Kroemer, G.) vol. 965 1–13 (Humana Press, 2013).

7. Basisty, N. et al. A proteomic atlas of senescence-associated secretomes for aging biomarker development. PLOS Biol. 18, e3000599 (2020).

8. Matjusaitis, M., Chin, G., Sarnoski, E. A. & Stolzing, A. Biomarkers to identify and isolate senescent cells. Ageing Res. Rev. 29, 1–12 (2016).

9. Lee, S. & Schmitt, C. A. The dynamic nature of senescence in cancer. Nat. Cell Biol. 21, 94–101 (2019).

10. Campisi, J. Cellular senescence: putting the paradoxes in perspective. Curr. Opin. Genet. Dev. 21, 107–112 (2011).

11. Gorgoulis, V. et al. Cellular Senescence: Defining a Path Forward. Cell 179, 813–827 (2019).

12. Mitsui, Y. & Schneider, E. L. Increased nuclear sizes in senescent human diploid fibroblast cultures. Exp. Cell Res. 100, 147–152 (1976).

13. Chen, J.-H. & Ozanne, S. E. Deep senescent human fibroblasts show diminished DNA damage foci but retain checkpoint capacity to oxidative stress. FEBS Lett. 580, 6669–6673 (2006).

14. Kusumoto, D. et al. Anti-senescent drug screening by deep learning-based morphology senescence scoring. Nat. Commun. 12, 257 (2021).

15. Collado, M. & Serrano, M. Senescence in tumours: evidence from mice and humans. Nat. Rev. Cancer 10, 51–57 (2010).

16. Campisi, J. CANCER: Suppressing Cancer: The Importance of Being Senescent. Science 309, 886–887 (2005).

17. Goldman, R. D. et al. Accumulation of mutant lamin A causes progressive changes in nuclear architecture in Hutchinson–Gilford progeria syndrome. Proc. Natl. Acad. Sci. 101, 8963–8968 (2004).

18. Martins, F., Sousa, J., Pereira, C. D., Cruze Silva, O.A.B. & Rebelo, S. Nuclear envelope dysfunction and its contribution to the aging process. Aging Cell 19, (2020).

19. Kassani, S. H., Kassani, P. H., Wesolowski, M. J., Schneider, K. A. & Deters, R. Breast Cancer Diagnosis with Transfer Learning and Global Pooling. in 2019 International Conference on Information and Communication Technology Convergence (ICTC) 519–524 (IEEE, 2019). doi:10.1109/ICTC46691.2019.8939878.

20. Tomita, H. et al. Deep Learning for the Preoperative Diagnosis of Metastatic Cervical Lymph Nodes on Contrast-Enhanced Computed ToMography in Patients with Oral Squamous Cell Carcinoma. Cancers 13, 600 (2021).

21. Gal, Y. & Ghahramani, Z. Dropout as a Bayesian Approximation: Representing Model Uncertainty in Deep Learning. ArXiv150602142 Cs Stat (2016).

22. Wen, Y., Vicol, P., Ba, J., Tran, D. & Grosse, R. Flipout: Efficient Pseudo-Independent Weight Perturbations on Mini-Batches. ArXiv180304386 Cs Stat (2018).

23. Fort, S., Hu, H. & Lakshminarayanan, B. Deep Ensembles: A Loss Landscape Perspective. ArXiv191202757 Cs Stat (2020).

24. Hewitt, G. et al. Telomeres are favoured targets of a persistent DNA damage response in ageing and stress-induced senescence. Nat. Commun. 3, 708 (2012).

25. Hernandez-Segura, A., Nehme, J. & Demaria, M. Hallmarks of Cellular Senescence. Trends Cell Biol. 28, 436–453 (2018).

26. Petr, M. A., Tulika, T., Carmona-Marin, L. M. & Scheibye-Knudsen, M. Protecting the Aging Genome. Trends Cell Biol. 30, 117–132 (2020).

27. Keijzers, G., Bakula, D. & Scheibye-Knudsen, M. Monogenic Diseases of DNA Repair. N. Engl. J. Med. 377, 1868–1876 (2017).

28. Idda, M. L. et al. Survey of senescent cell markers with age in human tissues. Aging 12, 4052–4066 (2020).

29. Collado, M., Blasco, M. A. & Serrano, M. Cellular Senescence in Cancer and Aging. Cell 130, 223–233 (2007).

30. Liggett, W. H. & Sidransky, D. Role of the p16 tumor suppressor gene in cancer. J. Clin. Oncol. 16, 1197–1206 (1998).

31. Campisi, J., Andersen, J. K., Kapahi, P. & Melov, S. Cellular senescence: A link between cancer and age-related degenerative disease? Semin. Cancer Biol. S1044579X11000502 (2011) doi:10.1016/j.semcancer.2011.09.001.

32. Burd, C. E. et al. Monitoring Tumorigenesis and Senescence In Vivo with a p16INK4a-Luciferase Model. Cell 152, 340–351 (2013).

33. Wang, B., Kohli, J. & Demaria, M. Senescent Cells in Cancer Therapy: Friends or Foes? Trends Cancer 6, 838–857 (2020).

34. Pathak, R. U., Soujanya, M. & Mishra, R. K. Deterioration of nuclear morphology and architecture: A hallmark of senescence and aging. Ageing Res. Rev. 67, 101264 (2021).

35. Filippi-Chiela, E. C. et al. Nuclear Morphometric Analysis (NMA): Screening of Senescence, Apoptosis and Nuclear Irregularities. PLoS ONE 7, e42522 (2012).

36. Neri, F., Basisty, N., Desprez, P.-Y., Campisi, J. & Schilling, B. Quantitative Proteomic Analysis of the Senescence-Associated Secretory Phenotype by Data-Independent Acquisition. Curr. Protoc. 1, e32 (2021).

37. Ronneberger, O., Fischer, P. & Brox, T. U-Net: Convolutional Networks for Biomedical Image Segmentation. in Medical Image Computing and Computer-Assisted Intervention – MICCAI 2015 (eds. Navab, N., Hornegger, J., Wells, W.M. & Frangi, A.F.) vol. 9351 234– 241 (Springer International Publishing, 2015).

