## Supplementary information for "Deep Learning Shows Cellular Senescence Is a Barrier to Cancer Development"

The supplementary information contains:

- Supplementary figure legends.
- Supplementary figure S1-S5.

**Supplementary figure legends**

**Figure S1** ***In vitro* models of senescence. a** Schematic of Ionizing radiation induced senescence. **b** Representative growth curve of cells undergoing replicative senescence. **c** Representative micrographs of ß-galactosidase staining in control, IR and RS cells.

**Figure S2 Performance characteristics of Several Models. a** Accuracy of raw model, tested on independent cell lines. **b** ROC curve of the raw model for independent cell lines. **C** Precision/Recall curve. **d** Accuracy of three-state senescent type detector model. **e** ROC curve with RS and IR for the type model. **f** Accuracy of normalized model. **g** ROC curve for the normalized model. **h** Accuracy of three-state senescent type detector model with normalized samples. **i** ROC curve for the type model with normalized samples. **j** Accuracy of Xception-based Bayesian neural network. **k** ROC curve for the Xception BNN. **l** Accuracy of InceptionV3-based Bayesian neural network. **m** ROC curve for the InceptionV3 BNN.

**Figure S3 Improving performance by adjusting thresholds. a** Histogram of predicted probabilities for Bayesian neural network. **b** Histogram of predicted probabilities for deep ensemble. **c** Accuracy and percent of samples evaluated with different classifier thresholds for single model. **d** Accuracy and percent of samples evaluated with different classifier thresholds for Xception-based Bayesian neural network. **e** Accuracy and percent of samples evaluated with different classifier thresholds for deep ensemble. **f** Accuracy and percent of samples evaluated with different classifier thresholds for deep ensemble with normalized samples. **g** Accuracy and percent of samples evaluated with different classifier thresholds for ensemble of RS-only for normalized samples. **h** Accuracy and percent of samples evaluated with different classifier thresholds for ensemble of IR-only for normalized samples.

**Figure S4 DNA Damage Foci and Morphological Characteristics of Predicted Senescence. a** Representative immunohistochemistry micrographs of nuclei with DNA damage foci staining. **b** 2D histogram (density heatmap) of predicted senescence and foci count per senescence type. **c** 2D histogram (density heatmap) of predicted senescence and foci count for premature aging diseases. **d** 2D histogram (density heatmap) of predicted senescence and foci count for murine astrocytes. **e** Predicted probability of RS senescence (n=5). **f** Predicted probability of IR senescence (n=5).

**Figure S5 Senescence metrics in human dermal fibroblasts. a** Predicted probability of RS senescence (n=169). **b** Predicted probability of IR senescence (n=169). **c** Correlation coefficiency matrix of metrics from nuclei. **d** 2D histograms (density heatmaps) of nuclei from the histology slides.

Figure S1

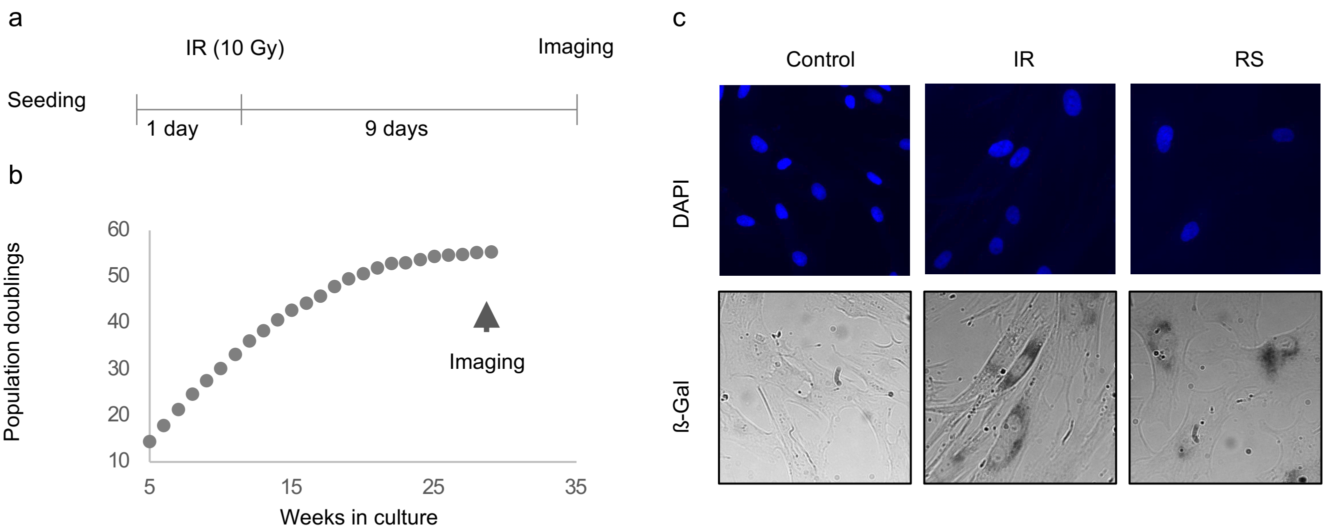
a
IR (10 Gy)
Imaging
Seeding
1 day
9 days
b
Population doublings
60
50
40
30
20
10
5
15
25
35
Weeks in culture
Imaging
c
Control
IR
RS
DAPI
ß-Gal

Figure S2

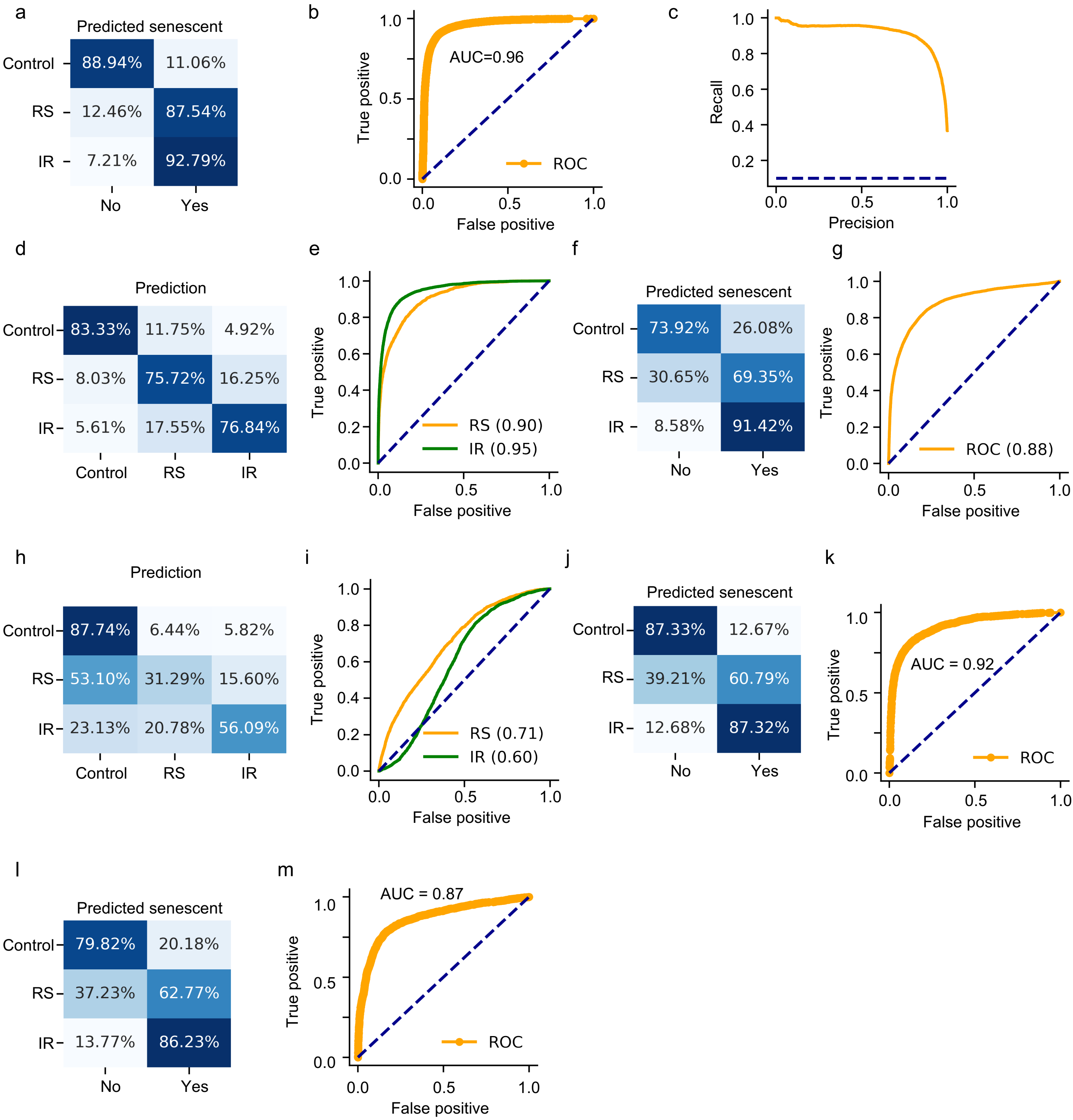
a
Predicted senescent
Control
RS
IR
88.94%
11.06%
12.46%
87.54%
7.21%
92.79%
No
Yes
b
AUC=0.96
ROC
True positive
False positive
c
Recall
Precision
d
Prediction
83.33%
11.75%
4.92%
8.03%
75.72%
16.25%
5.61%
17.55%
76.84%
e
RS (0.90)
IR (0.95)
f
73.92%
26.08%
30.65%
69.35%
8.58%
91.42%
g
ROC (0.88)
h
87.74%
6.44%
5.82%
53.10%
31.29%
15.60%
23.13%
20.78%
56.09%
i
RS (0.71)
IR (0.60)
j
87.33%
12.67%
39.21%
60.79%
12.68%
87.32%
k
AUC = 0.92
l
79.82%
20.18%
37.23%
62.77%
13.77%
86.23%
m
AUC = 0.87

Figure S3

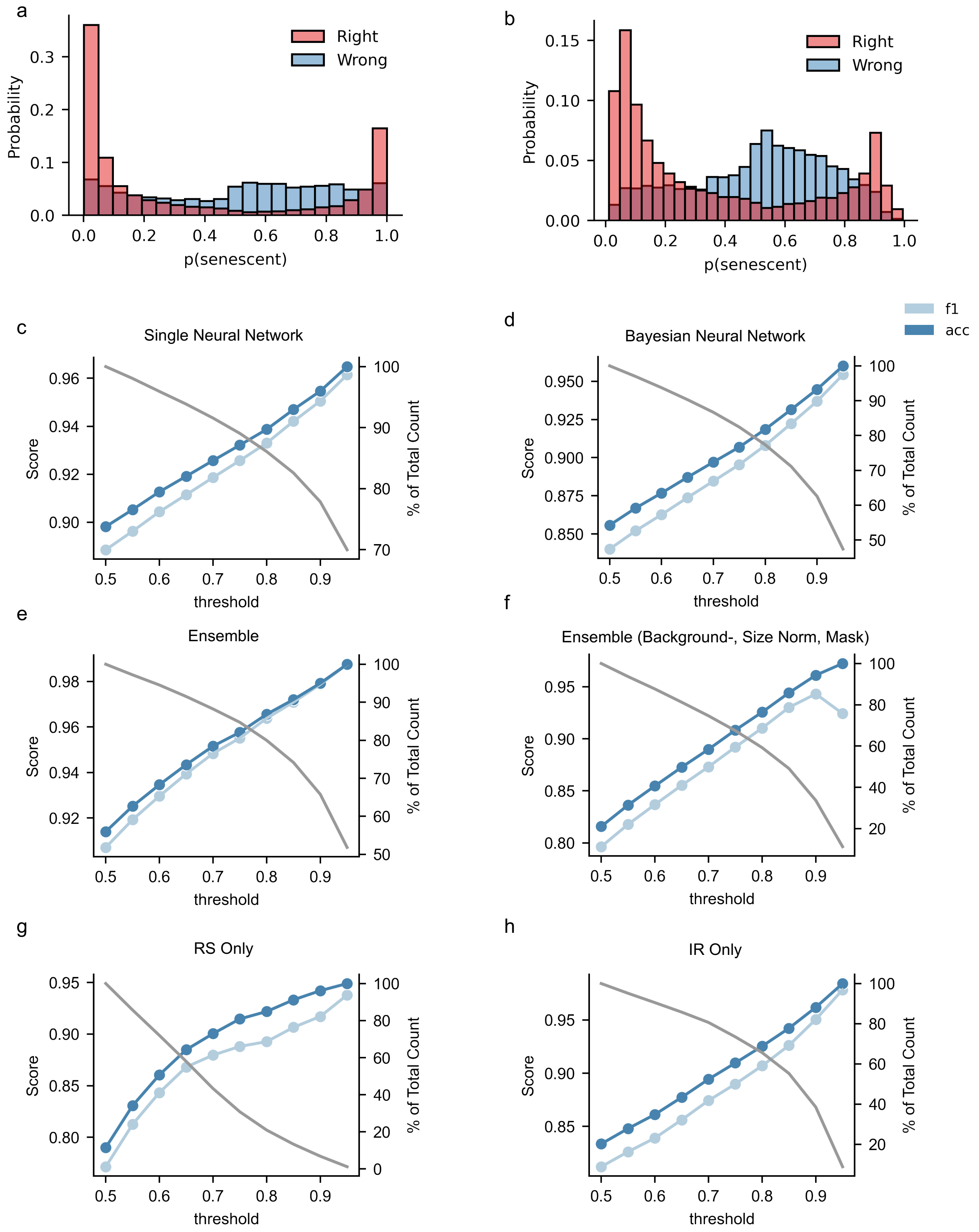
a
Right
Wrong
Probability
p(senescent)
b
Right
Wrong
Probability
p(senescent)
c
Single Neural Network
d
Bayesian Neural Network
f1
acc
e
Ensemble
f
Ensemble (Background-, Size Norm, Mask)
g
RS Only
h
IR Only
Score
% of Total Count
threshold

Figure S4

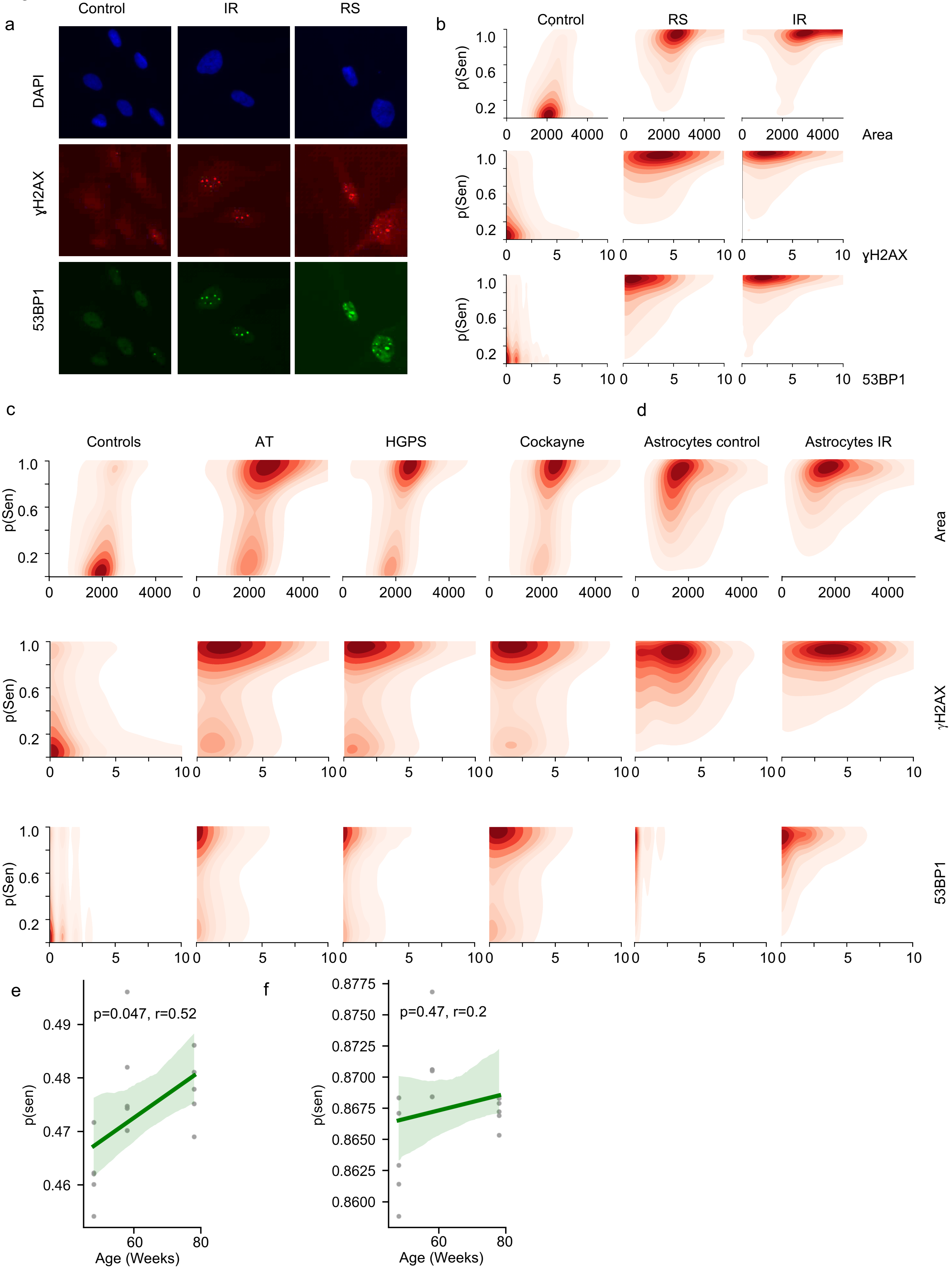
a
Control
IR
RS
DAPI
γH2AX
53BP1
b
Control
RS
IR
p(Sen)
1.0
0.6
0.2
0
2000
4000
Area
5
10
γH2AX
53BP1
c
Controls
AT
HGPS
Cockayne
d
Astrocytes control
Astrocytes IR
e
p=0.047, r=0.52
0.49
0.48
0.47
0.46
p(sen)
60
80
Age (Weeks)
f
0.8775
0.8750
0.8725
0.8700
0.8675
0.8650
0.8625
0.8600
p=0.47, r=0.2
p(sen)
60
80
Age (Weeks)

Figure S5

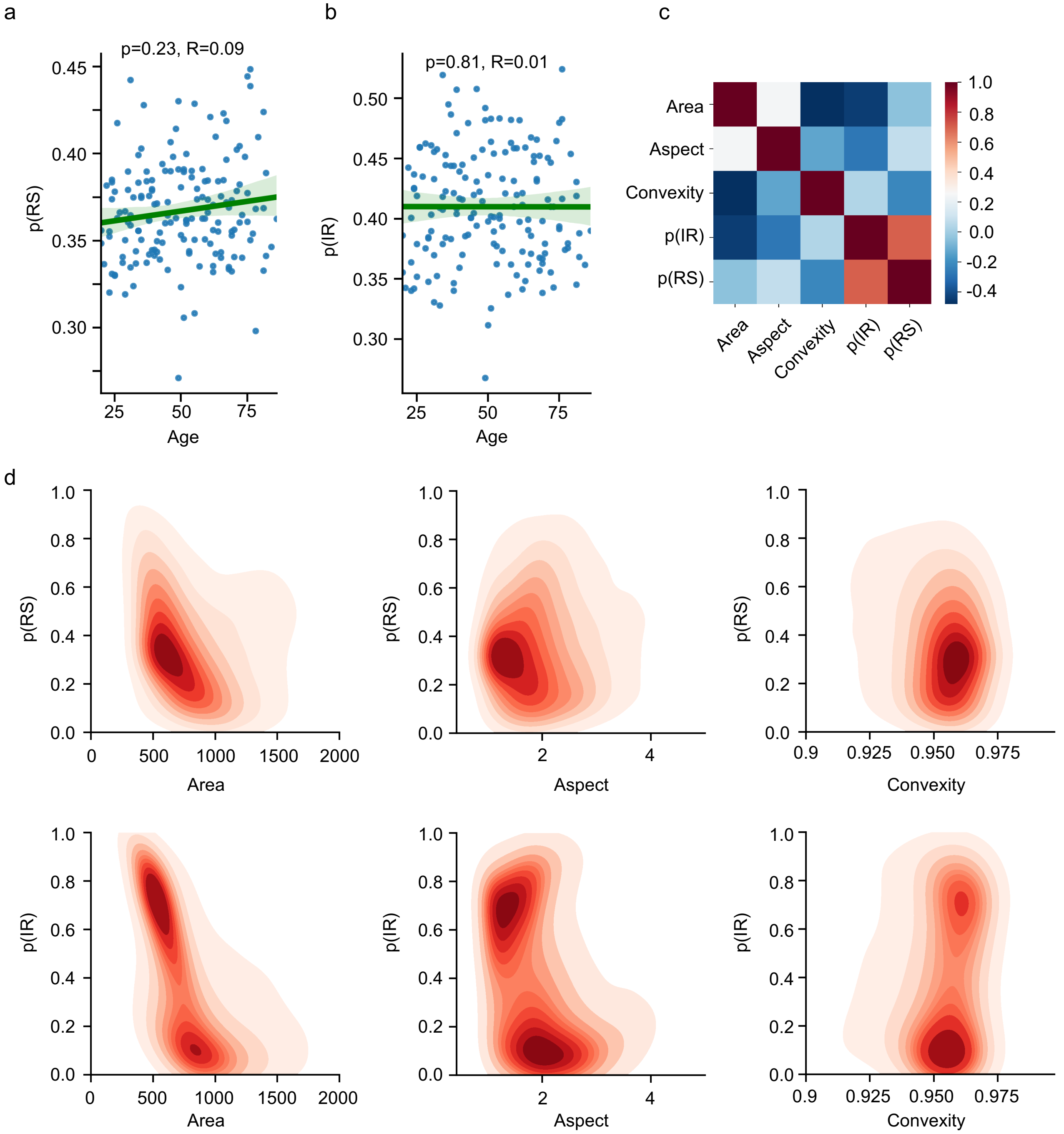
a
p=0.23, R=0.09
p(RS)
Age
b
p=0.81, R=0.01
p(IR)
Age
c
Area
Aspect
Convexity
p(IR)
p(RS)
d
p(RS)
p(IR)
Area
Aspect
Convexity
